# Solvent accessibility of chlorine-reactive amino acid residues in icosahedral virus structures: A meta-analysis

**DOI:** 10.64898/2026.06.17.732355

**Authors:** Chonglin Zhu, Kerry Prinsen, Leah Ward, Mira Chaplin, Meng Shen, Shotaro Torii, Kathryn Kauffman, Yinyin Ye

**Affiliations:** Department of Civil, Structural and Environmental Engineering, University at Buffalo, Buffalo, New York 14260, United States; Department of Civil and Environmental Engineering, University of Michigan, Ann Arbor, Michigan 48105, United States; Department of Physics, California State University, Fullerton, California 92831, United States; Department of Urban Engineering, School of Engineering, The University of Tokyo, Bunkyo City, Tokyo 113-8654, Japan; Department of Oral Biology, University at Buffalo, Buffalo, New York 14214, United States

**Keywords:** Chlorine disinfection, solvent-accessible areas, water treatment, structural analysis, structural bioinformatics, viruses

## Abstract

Free chlorine reacts with viral proteins, but the protein structural determinants of viral resistance to chlorine treatment remain poorly understood. Here, we curated a dataset of 498 icosahedral virus structures, including intact virions and virus-like particles (VLPs), from the Protein Data Bank. Surprisingly, only 6.6% of these structures are associated with published viral chlorine inactivation rate constants (*k*_obs_). In these matched cases representing 12 virus families, total and maximum solvent accessible surface areas (SASA) of methionine residues within viral attachment and entry proteins correlated significantly with median *k*_obs_ (Pearson’s *r* = 0.83 and 0.45, respectively; *p* < 0.05), suggesting a critical role of methionine exposure in viral resistance phenotypes. Across the full curated dataset, fuzzy c-means clustering upon total and maximum SASA profiles of chlorine-reactive residues demonstrates that the common surrogate panel (MS2, PhiX174, Phi6, PRD1, and PR772) fails to represent the SASA diversity of human viruses. Instead, VLPs and novel phages may serve as better surrogates for chlorine treatment due to SASA profile similarities to human viruses. Our findings highlight that residue SASA features provide a quantitative baseline for screening viral resistance to chlorine and offer a data-driven strategy to select structurally representative virus surrogates for future disinfection studies.

**SYNOPSIS:** Solvent accessibility of chlorine-reactive residues correlates strongly with viral chlorine resistance, providing a robust quantitative metric to compare chlorine resistance phenotypes across viral capsids.

**TOC:** 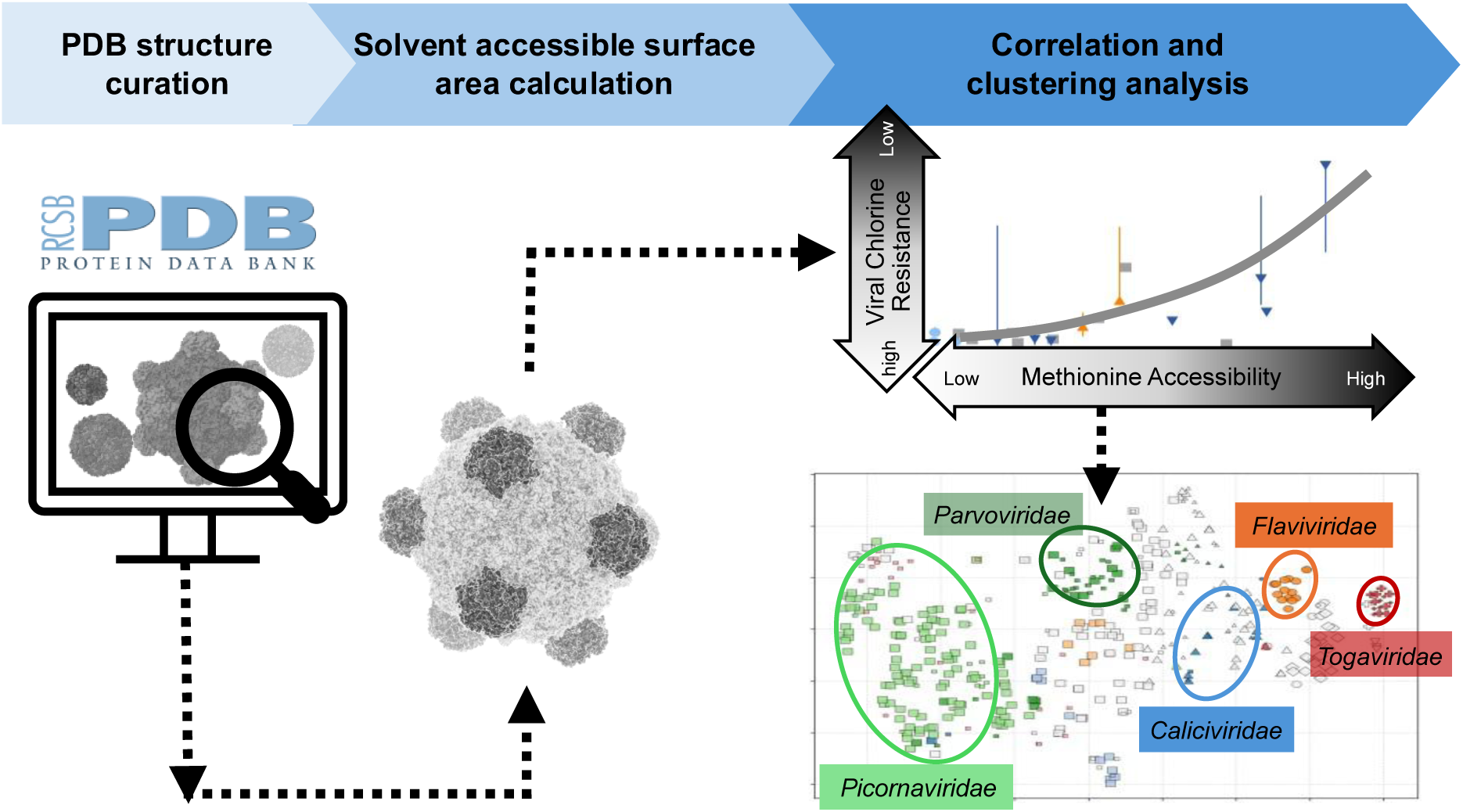

## INTRODUCTION

The inactivation kinetics of free chlorine disinfection of viruses vary substantially in the environment.^1, 2^ To ensure that disinfection approaches respond effectively to emerging viral threats, it is crucial to characterize the mechanisms underlying such variability. At the molecular level, reactions between free chlorine and viral proteins drive virus inactivation.^3^ In human astrovirus and norovirus, chlorine treatment leads to significant oxidative damage in viral capsids with the formation of carbonyl products,^4^ while the genome remains largely intact. Chlorine inactivation studies of Phi6, PhiX174, and T4 bacteriophages have reported correlations between the reaction kinetics of specific methionine-containing peptides and virus inactivation.^5, 6^ Additionally, peptides in MS2 capsid decay at varying rates during chlorine treatment, and the cumulative peptide decay accounts for approximately 50% of the loss in genome injection function.^7^

Within a bottom-up framework for predicting viral protein reactivity from their building blocks, a specific subset of amino acid residues is reactive with free chlorine. Pattison *et al.*,^8^ utilized UV spectrometry and quantified the absolute reactivity of free amino acids with free chlorine, with the most reactive methionine (3.8 × 10^7^ M^−1^ s^−1^) ≈ cysteine (3.0 × 10^7^ M^−1^ s^−1^), followed by histidine (1.0 × 10^5^ M^−1^ s^−1^) > tryptophan (1.1 × 10^4^ M^−1^ s^−1^) > lysine (5.0 × 10^3^ M^−1^ s^−1^) >> tyrosine (44 M^−1^ s^−1^) > arginine (26 M^−1^ s^−1^). Although free amino acid reaction rates serve as a useful baseline, relative reactivities shift when these residues are incorporated into complex viral proteins. The half-life monitoring of residues in the MS2 capsid with free chlorine reported a modified reactivity: methionine > tryptophan > tyrosine > lysine.^9^ The discrepancies between the relative reactivities of free and protein-bound tyrosine and lysine residues can be attributed to intramacromolecular chlorine transfer between lysine chloramine and nearby lysine residues.^9^ In addition, protein folding structures also regulate amino acid reactivity with chlorine.^7, 10^ Recent quantitative proteomic results demonstrate that peptides containing buried methionine residues remain shielded from chlorine attack compared to the peptides containing exposed methionine residues.^5, 6^ Together, these findings highlight that both the primary sequence and geometric layout are critical factors affecting viral protein reactivity.

The reactivity of amino acids or peptides in folded proteins is related to their solvent accessibility. Indeed, the reaction kinetics of lysine residues with 1,2-cyclohexanedione correlate directly with the solvent accessible surface areas (SASA) derived from high-resolution X-ray structures.^11^ Similarly, in the singlet oxygen-mediated oxidation of glyceraldehyde-3-phosphate dehydrogenase, the oxidation rates of histidine residues are governed by the residue exposure.^12^ This relationship has been extended to free chlorine reacting with bacteriophages, where the decay rates of methionine-containing peptides within the phage capsids are linked to the SASA of methionine residues and the average exposure of peptides.^5^ However, it remains unclear whether the capsid reactivity observed in phages accurately represents that of human or other emerging viruses. To investigate the structural diversity of viral capsids, the Protein Data Bank (PDB) serves as a comprehensive repository of atomic-resolution capsid structures resolved via cryo-electron microscopy and X-ray crystallography across different virus families.^13^ These PDB structures have yet to be systematically analyzed in the context of disinfection and chemical susceptibility.

In this study, we screened icosahedral virus capsid structures from PDB to curate a dataset of intact virions and virus-like particles (VLPs) suitable for modeling viral structures during water treatment processes. We developed a computational pipeline to calculate the SASA profiles of chlorine-reactive residues for assembled capsid structures. The resulting SASA values were leveraged to: (1) identify structural features correlated with viral chlorine resistance by benchmarking SASA values of chlorine-reactive residues against known chlorine inactivation rate constants of viruses, and (2) assess structural similarities of bacteriophage surrogates and human pathogens through machine learning-based clustering analysis. Our SASA-based framework for capsid structural analysis provides a quantitative structural component that complements current genome-oriented models for virus inactivation kinetics prediction, offering a more holistic approach to understanding viral resistance to oxidative disinfectants.

## MATERIALS AND METHODS

### Extraction of viral icosahedral virus structures from PDB

Icosahedral virus structures were retrieved in the Research Collaboratory for Structural Bioinformatics Protein Data Bank (RCSB PDB)^14^ on January 26, 2026. To ensure a comprehensive extraction of high-resolution (<= 5 Å), experimentally determined capsid structures associated with viruses or phages, we deployed two complementary attribute searches (**Figure S1**): Search A for perfect icosahedral symmetry (Symmetry Type = “Icosahedral” AND Symmetry Symbol = “I” AND Refinement Resolution <= 5 Å AND (Experimental Method = “X-RAY DIFFRACTION” OR Experimental Method = “ELECTRON MICROSCOPY”) AND (Structure Title HAS ANY OF WORDS “phage” OR Structure Title HAS ANY OF WORDS “virus” OR Text search inside all attributes = “capsid”)), while Search B to capture icosahedral capsids containing a single maturation protein (Symmetry Type = “Cyclic” AND (Symmetry Symbol = “C1” OR Symmetry Symbol = “C5”) AND Refinement Resolution <= 5 Å AND (Experimental Method = “X-RAY DIFFRACTION” OR Experimental Method = “ELECTRON MICROSCOPY”) AND (Structure Title HAS ANY OF WORDS “phage” OR Structure Title HAS ANY OF WORDS “virus” OR Text search inside all attributes = “capsid”)). Consequently, 1418 and 1266 structures were exported from Search A and B, respectively, into an Excel sheet, along with the structural information, including PDB ID, PubMed ID (if any), refinement resolution, and structure title. The initial screening was performed by removing structures whose titles indicated engineered protein cages, virus-antibody complexes, immature capsids, expanded capsids, viruses treated by or complexed with anti-viral drugs, viral vectors (i.e., adeno-associated viruses), or other non-viral structures. The remaining 884 structures were further manually filtered based on the structure description on the PDB website and a full-text review of the publication of the structure to remove structures determined in extreme pH (pH < 5 or pH > 10), naturally occurring empty capsids, inactivated viruses, and engineered mutants used for mechanistic studies of virus-host interactions. This process yielded 498 final structures consisting of infectious virus particles (i.e., virions) and VLPs for subsequent analysis (**Table S1**).

### Cross-reference of structure-relevant information

For each selected structure, the PDB ID was used to retrieve structural information from the RCSB PDB database via a GraphQL-based application programming interface (API),^15^ including molecule names, chain IDs, and associated UniProt IDs of the structural proteins. To cross-reference these structure-associated proteins in UniProtKB, including protein and gene names, the UniProt REST API was used, with responses parsed into JavaScript Object Notation (JSON).^16^ In cases where a UniProt ID was unavailable, the protein name was manually assigned based on the publication of the structure or the PDB molecule names. Viral names and strains associated with the PDB structure were extracted from the publication or searched in the UniProtKB for the organism and strain names using the protein UniProt ID. Virus lineage (realm, class, order, family, genus, species) was determined according to the International Committee on Taxonomy of Viruses (ICTV) Virus Metadata Resource (VMR_MSL40.v2.20260223).^17^ Comprehensive metadata mentioned above were provided in **Table S1**. Furthermore, the common hosts of viruses, viral genome structures, enveloped/nonenveloped status, virion proteins, and a list of specific proteins responsible for viral attachment and entry were manually retrieved from the Virus-Host DB^18^ and ViralZone^19^ for each virus family and summarized in **Table S2**.

### Solvent accessible surface area (SASA) of residues within structural proteins

Biological assemblies of viral capsid structures (assembly ID = 1) were downloaded from the PDB archive^20^ as compressed mmCIF format. The compressed structures were then parsed using the Bio.PDB.MMCIFParser module in Biopython (v1.85),^21^ and were subsequently cleaned up in two steps using the cmd module in PyMOL conda packages (v3.1.7.2):^22^ (i) the removal of all water molecules, and then (ii) the removal of non-amino acid ligands and heteroatoms. After the cleanup, the Biopython Bio.PDB.SASA module was used to calculate solvent accessible surface areas (SASA) at the amino acid residue level based on the rolling ball algorithm developed by Shrake & Rupley^23^ with a default radius of a water molecule (probe radius) as 1.40 Å and surface resolution (n points) as 300. For a given protein m in the structure S, the SASA of residue A at residue position *i* in copy *j*, *S*_*mi,j*_, was first determined. Subsequently, these values were used to quantify the total SASA of residue A in protein m 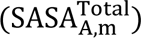 and the maximum SASA of residue A 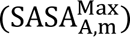 in protein m:

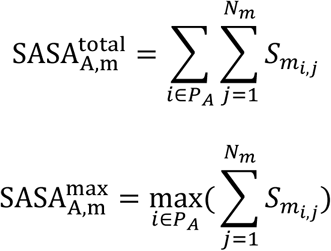

Where *N_m_* is the copy number of protein m in the capsid; *P*_*A*_ contains all position indices *i* where the residue is A, *P*_*A*_ = {*i*: residue in protein m is of type A}. The 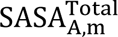 and 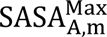 values of each protein m within a given PDB structure in the curated dataset were calculated for all 20 common amino acids and summarized in **Table S3**.

To test the impact of calculation workflow on the outcome SASA values, PDB structures (2BPA, 8AT5, 5C4W, and 7BG6) were selected to evaluate the cleanup strategies (*i.e.*, no cleanup, water removal, and all cleanup), probe radius (*i.e.*, 1.20, 1.40, 1.60, and 2.20 Å), and surface area resolution n points (*i.e.*, 100, 300, and 1000). Details of the analysis design were provided in **Table S4.**

### Mapping protein SASA to virus inactivation rate constants by free chlorine

The observed virus inactivation rate constants by free chlorine (*k_obs_*, L mg^−1^ min^−1^) were curated from two sources: (i) a comprehensive systematic review published previously^1^ and (ii) recent experimental results not captured in the review.^5, 24^ Firstly, viruses with the rate constants summarized in the review^1^ were compared with the virus information associated with the PDB structures in **Table S1** to match at the strain level. To maintain environmental conditions relevance for structures, the rate constants of the matched strains were refined using the following criteria: (a) the kinetics data were limited to buffered suspensions (calcium, carbonate, phosphate, phosphate carbonate, and phosphate saline) and treated waters (treated water, treated groundwater, treated surface water, and tap water), (b) pH values were constrained to a range of 5.1 to 9.9 consistent with the pH range used for screening the PDB structures, and (c) temperatures constrained between 4 to 30 °C. Secondly, these refined rate constants were supplemented with rate constants for a mutant of coxsackievirus B5 strain Faulkner,^24^ bacteriophages (T4, PhiX17, MS2),^5^ and bacteriophage Phi6 (**Figure S2**) measured in phosphate buffer (pH 7.0-7.5) at room temperature. Given that the rate constants followed skewed distributions for most of the virus strains, the median *k_obs_*, and the 25^th^ and 75^th^ percentiles (Q1 and Q3) were calculated (**Table S5**).

To build a mechanistic link between capsid structures and virus chlorine resistance, seven chlorine-reactive residues were included, *i.e.*, methionine (Met, M), cysteine (Cys, C), histidine (His, H), tryptophan (Trp, W), lysine (Lys, K), tyrosine (Tyr, Y), and arginine (Arg, R). The SASA of these residues was calculated for specific protein groups based on whether they are responsible for viral attachment and entry. For each virus structure, the total SASA of residues in protein group G_k_ 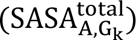 and the maximum SASA of residues in protein group G_k_ 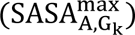 were derived as follows:

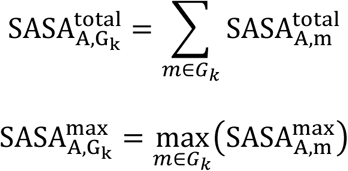

Where G_k_, k ɛ{1, 2} is the protein group. Specifically, G_1_ contains viral attachment and entry proteins, and G_2_ includes the rest of the proteins in the structure. If one viral strain is associated with multiple PDB structures, their average 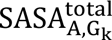 and 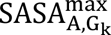 values were used for correlation analysis (**Table S5**).

### Statistical analysis

To evaluate relationships between residue SASA values calculated at various probe radii and surface resolutions, Pearson correlation analysis was performed. Results with Pearson’s correlation coefficient *r* > 0.9 and p-value < 0.05 were considered as highly positive correlations.

For viral strains with known capsid structures and chlorine inactivation kinetics, we employed Pearson’s correlation analysis to evaluate the relationships between median viral chlorine resistance (*k_obs_*) and each of the total and maximum SASA 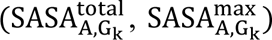 of seven chlorine-reactive amino acids within protein groups G_1_ and G_2_. To assess if the Pearson’s correlation coefficients were disproportionally driven by outliers with the small dataset, non-parametric Spearman’s rank and Kendall’s rank correlation analyses were conducted in parallel. A substantial drop in statistical significance during non-parametric testing was used to flag potential outlier distortion. SASA variables that correlated significantly with the median *k_obs_* (*p*-value < 0.05) across the three correlation methods were then selected for subsequent linear regression analysis in GraphPad Prism (v10.6.0, GraphPad Software). To confirm that predictor variables were independent, multicollinearity was assessed using the variance inflation factor^25^ and pairwise Pearson correlation analysis.

To compare the total and maximum SASA profiles of seven reactive amino acid residues within the group of viral attachment and entry proteins, fuzzy c-means clustering^26^ was applied, which primarily grouped structures based on their absolute SASA values in our case. The number of clustering groups was adjusted to ensure that structures from the same virus families were mostly represented in one cluster. Subsequently, clusters were visualized in a t-distributed Stochastic Neighbor Embedding (t-SNE) plot,^27^ where nearby structures would have higher similarities in their residue SASA profiles than distant structures.

### Code availability

The Python scripts for metadata retrieval via PDB GraphQL and UniProt REST APIs, PDB structure cleanup, residue-level SASA calculation, and example data are available on GitHub at https://github.com/MarcoPineapollo/Virus-structural-analysis-code.

## RESULTS AND DISCUSSION

### Overview of curated PDB structure dataset

The curated dataset comprised 498 icosahedral viral capsids, including 119 VLPs and 377 virion structures (**Table S1**). While VLPs represent non-infectious virus particles whose capsid composition may diverge from wild-type counterparts,^28^ they are indispensable for understanding the capsid structures of nonculturable or difficult-to-culture viruses such as human norovirus.^29^ We therefore stratified our data by analyzing VLP and virion structures as two independent cohorts in the subsequent investigation. Within the dataset, 473 structures are associated with classified viruses, primarily originating from four major realms, *i.e.*, *Riboviria*, *Duplodnaviria*, *Floreoviria*, and *Varidnaviria*, and representing six Baltimore classes,^30^ with (+)ssRNA viruses being the most prevalent (**Figure 1A**). Although the dataset encompasses 71 virus families, the overall distribution is heavily skewed. The top 10 families comprise approximately 55% of the structures, led by *Picornaviridae* (n = 121) and *Parvoviridae* (n = 45) (**Figure 1B**; **Table S2**). These highly represented virus families are complemented by a wide array of tailed bacteriophages, archaeal phages, virophages, and plant viruses, with most families contributing fewer than four structures (**Table S2**). Furthermore, non-enveloped viruses, including those with internal lipids, are dominant in the dataset (**Figure 1B**). Note that our search was restricted to icosahedral structures, filamentous viruses (*e.g.*, Ebola virus), rod-shaped viruses (*e.g.*, pepper mild mottle virus), pleomorphic viruses (*e.g.*, influenza virus^31^ and coronavirus^32^), and tail components of tailed phages were excluded.

**Figure 1.**
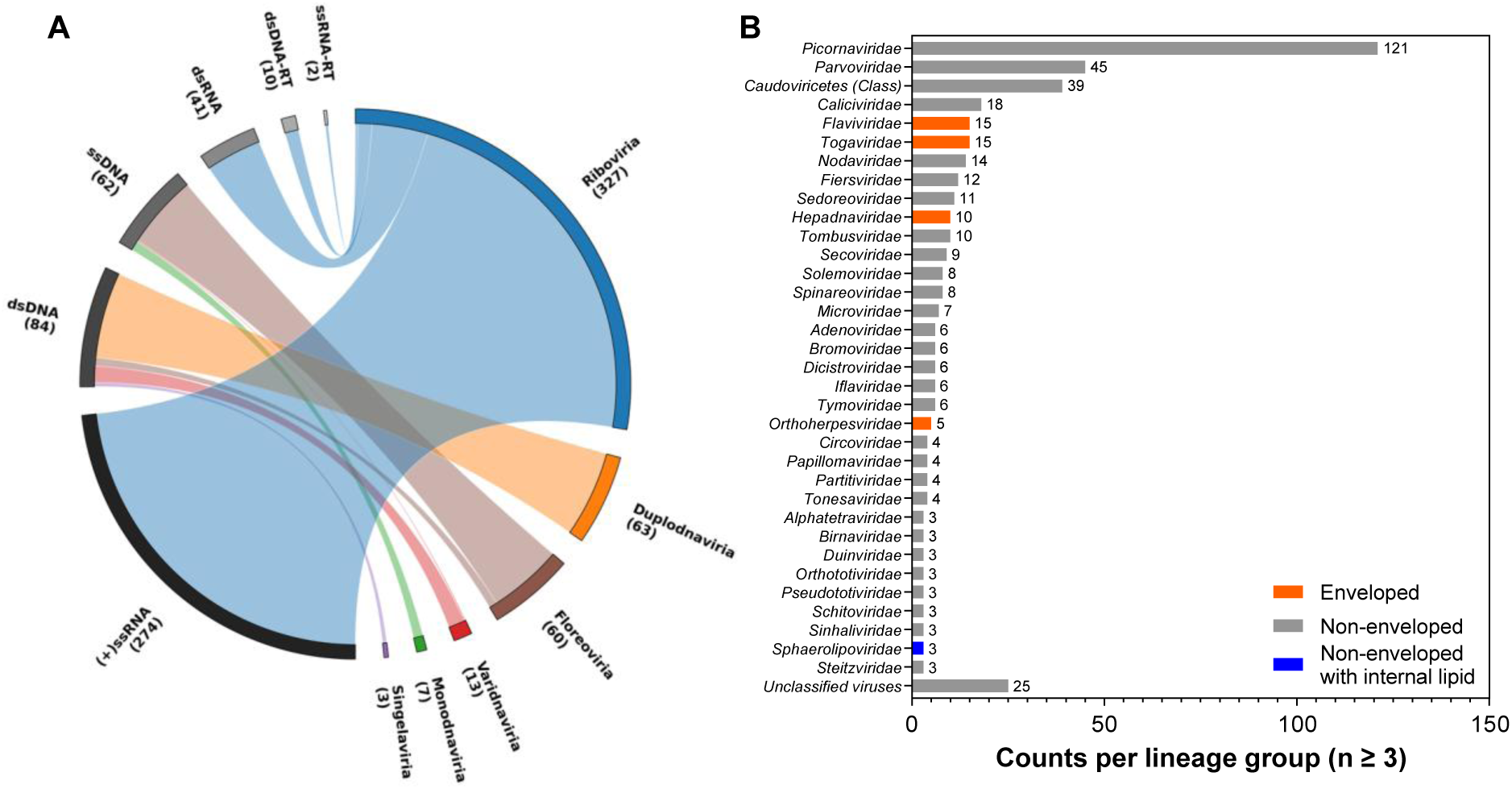
Overview of the curated PDB dataset for icosahedral virus structures. (A) Connections between the Baltimore classes and virus realms. The thickness of each ribbon is proportional to the number of structures within that specific Baltimore class-realm pair. (B) Structural distribution of taxonomic lineages at the family or class level with three or more structures. Bar colors denote the envelope status, classified as enveloped, non-enveloped, or non-enveloped with an internal lipid layer. Viruses in the group of *Caudoviricetes* (Class) lack an assigned family, and unclassified viruses lack assigned class, order, and family.

### SASA calculation validation for viral capsids

We first systematically evaluated the impact of the SASA calculation parameters, including structure cleanup strategies, probe radii, and surface area resolution, on the output SASA values using virion structures of coxsackieviruses (8AT5, 5C4W), rhinovirus (7BG6), and phage PhiX174 (2BPA). These structures were selected because they accounted for various complexities, including the presence of water molecules, host-derived lipid molecules bound within the capsid pockets (e.g., sphingosine, palmitic acid, and myristic acid), and partially resolved nucleic acid structure (**Table S4**). Our analysis demonstrated that removing water molecules yielded significantly higher residue SASA values at many sites, whereas the removal of pocket factors or nucleic acids only slightly exposed amino acids at a few specific sites that were otherwise buried by these molecules (**Figure S3A**). Because pocket factors and viral nucleic acids were not universally resolved or present across all viral structures, we computationally removed all non-amino acid molecules prior to SASA calculations as a consistent cleanup strategy.

We next examined the influence of probe radius, which is defined as the radius of a spherical solvent molecule used to probe the surface of amino acid residues. While a 1.4 Å radius is the default setting for a water molecule, a previous study used a 1.2 Å probe to estimate the SASA of residues accessible by singlet oxygen.^12^ In chlorine treatment, the concentrations of reactive species are pH-dependent, comprising hypochlorous acid (HOCl), hypochlorite ions (OCl^−^), aqueous chlorine (Cl_2_), and chloride ions (Cl^−^),^33^ and thus determining a single representative radius for chlorine treatment would be challenging. Instead of identifying one radius value for our analysis, we investigated how the deviation from the default water probe size influences the SASA values. Our results showed that absolute SASA values were sensitive to the probe radius. Small probes penetrated deeper into the structure surfaces, resulting in higher SASA values compared to large probes (**Figure S3B**). In particular, the probe at 2.2 Å was inaccessible to more residues than the probe at 1.2 Å. Despite the values change, SASA values calculated at non-default radius settings (probe radii of 1.2, 1.6, and 2.2 Å) remained highly correlated with the SASA values at the default setting (all Pearson’s r > 0.93, p<0.001; **Table S6**). These findings suggest that the default probe size is valid for investigating the correlations between residue SASA and virus resistance to free chlorine. The probe size would become primarily critical for quantitative applications where buried residues significantly contribute to predictive values, such as molecular solubility.^34^

The resolution of the surface area is approximated by a set of test points (n), where the SASA is calculated by rolling the spherical probe over the target surface. A higher number of test points yields more accurate estimations, but significantly increases computational demand.^35^ We therefore profiled the deviations of SASA values at resolutions of n = 100 and 300 from the n = 1000 setting as the high-accuracy baseline. The results showed that SASA values calculated at n = 100 were more dispersed than those at n = 300 when compared to the baseline (**Figure S3C**), but the resulting SASA values were all highly correlated with the baseline values (Pearson’s r > 0.99, p < 0.001; **Table S6**). Specifically, the n = 100 estimates frequently produced errors exceeding ±20 Å^2^, whereas the n = 300 estimates were more tightly clustered within a ±20 Å^2^ range (**Figure S3C**). To balance the computational throughput with numerical precision, we selected n = 300 as the standard resolution for all structure calculations.

In summary, we implemented and validated a standardized workflow for SASA calculations, utilizing consistent cleanup steps, a probe radius of r = 1.4 Å, and a surface area resolution of n = 300. The adjustment of probe radius and resolution values resulted in highly correlated SASA values, suggesting that the selection of these two parameters would not affect the conclusions for correlation-based analyses.

### Correlation between residue SASA and viral chlorine resistance across the selected viral strains

Twenty-three viral strains associated with previously reported chlorine inactivation rate constants (*k_obs_*) and at least one virion structure in the curated PDB dataset were selected for analysis (**Table S5**). These strains belong to 12 virus families, with the highest representation from *Picornaviridae* (n = 8), *Fiersviridae* (n = 4), and *Tectiviridae* (n = 2). The remaining 9 families, namely *Adenoviridae*, *Caliciviridae*, *Cystoviridae*, *Demerecviridae*, *Microviridae*, *Parvoviridae*, *Polyomaviridae*, *Sedoreoviridae*, and *Straboviridae*, are each represented by a single strain.

To investigate whether the solvent accessibility of chlorine-reactive residues in viral capsids has a cumulative effect on viral resistance to free chlorine disinfection, we first calculated the total SASA for residues within the specific proteins responsible for attachment and entry (designated as protein group G1). We then applied Pearson correlation alongside non-parametric Spearman and Kendall rank correlation analyses to verify whether the linear trends captured by Pearson’s coefficient were driven by outliers within our small dataset. Among the seven reactive residues evaluated, only the total SASA of Met correlated significantly with the median virus inactivation rate constants for all three statistical methods (Pearson’s *r* = 0.83, Spearman’s *ρ* = 0.60, Kendall’s *τ* = 0.46, all *p* < 0.05; **Figure 2**; **Table S7**). This robust correlation aligns with previous studies on chlorine-treated MS2, PhiX174, T4, and Phi6, which found that Met-containing peptides in viral capsids decayed more rapidly than those without Met residues.^5, 6^ Despite free Cys residues being as highly reactive with free chlorine as Met,^8^ total Cys SASA showed insignificant correlation with virus inactivation kinetics (*r* = 0.16, *ρ* = 0.080, *τ* = 0.053, all *p* > 0.4; **Figure 2**; **Table S7**). This lack of correlation is explainable by the data points of Phi6 and enterovirus 70 strain J670/71. Both viruses lack accessible Cys residues in their capsids yet remain highly susceptible to free chlorine (**Table S5**). Furthermore, among the other capsids studied, the total Met SASA was, on average, 10-fold greater than the total Cys SASA (**Table S5**). This disparity is expected, given that Cys is natively the least abundant amino acid among the twenty common residues in a wide range of organisms,^36^ and that Cys residues behave hydrophobically in proteins.^37^

**Figure 2.**
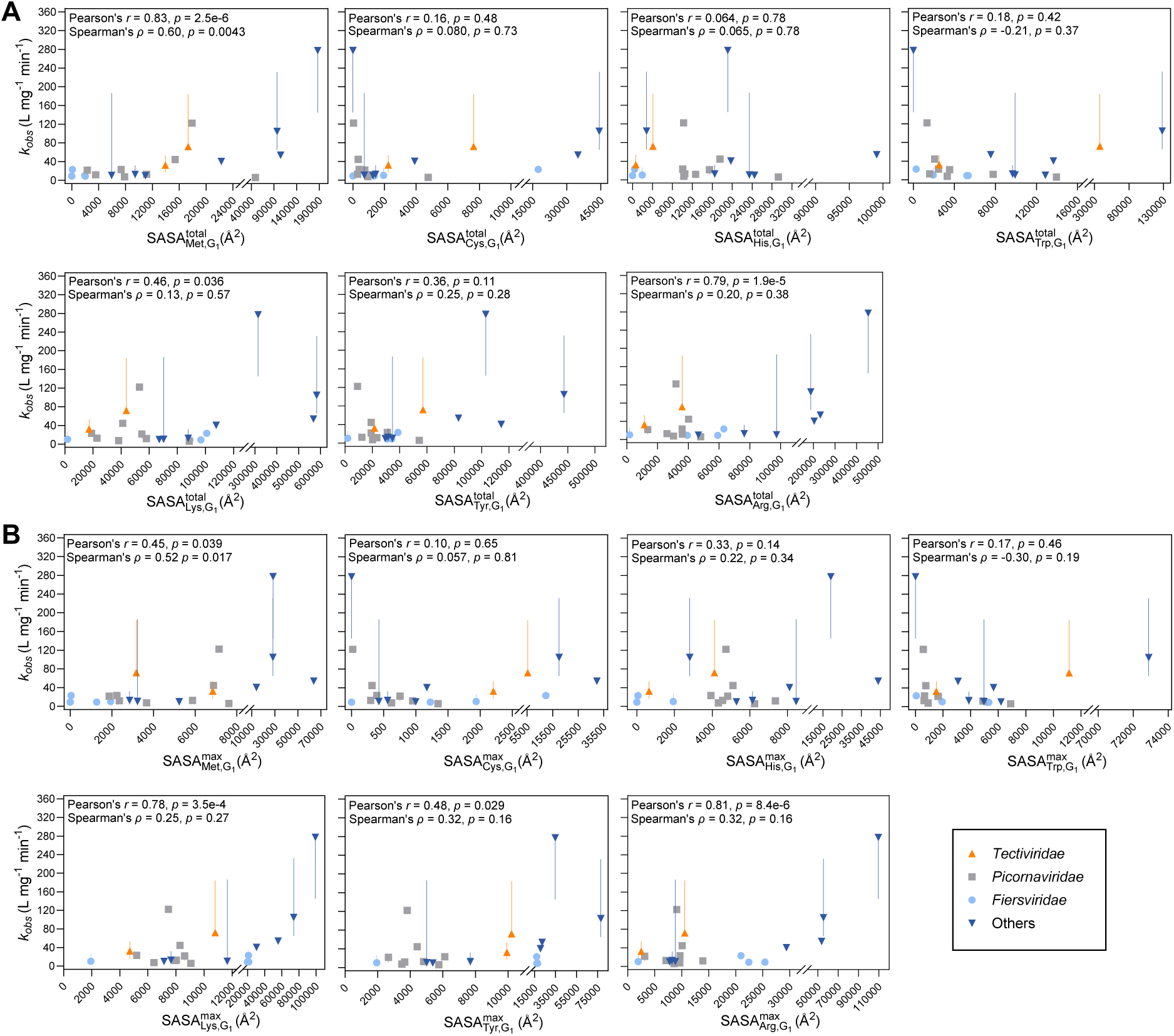
Correlations between virus inactivation rate constants by free chlorine (*k_obs_*) and (A) total solvent accessible surface areas (SASA) and (B) maximum SASA of chlorine-reactive residues within the protein group G_1_, directly involved in virus attachment and entry. The residues analyzed include methionine (Met), cysteine (Cys), histidine (His), tryptophan (Trp), lysine (Lys), tyrosine (Tyr), and arginine (Arg). Error bars represent the interquartile range (Q1 and Q3) of the median rate constants. Pearsons’s correlation coefficient (*r*) and *p-*values, and Spearman’s rank correlation coefficient (*ρ*) and *p*-values are displayed within each panel. Corresponding Kendall’s rank correlation coefficient (*τ*) and *p*-values were summarized in **Table S7**.

Beyond the total SASA as the cumulative exposure, we further evaluated whether viral chlorine resistance is governed by the accessibility of the most exposed reactive residues in the protein group G_1_. To this end, we calculated the maximum SASA for each type of chlorine-reactive residue to quantify the most exposed individual amino acid. The results showed that the maximum Met SASA was correlated significantly with virus inactivation rate constants (*r* = 0.45, *ρ* = 0.52, *τ* = 0.37, all *p* < 0.05; **Figure 2**; **Table S7**), suggesting that reactions in the most exposed Met residue may lead to virus inactivation. Consistently, previous studies of PhiX174, T4, and Phi6 reported that the decay kinetics of the most reactive Met-containing peptides, specifically those located in proteins responsible for attachment and entry, closely followed virus inactivation kinetics.^5, 6^

In contrast to the correlations observed for attachment and entry proteins, no significant correlations were observed between the inactivation rate constants and either the total or maximum SASA of reactive residue in the protein group G_2_, which is not directly responsible for attachment and entry functions (**Table S7**). Similar finding were previously reported in T4 phage, whose overall chlorine resistance was more accurately explained by the reactivity of proteins within the tail fibers, which are responsible for host attachment, rather than the proteins within its head capsid.^5^

Building upon the above correlation analyses, we identified moderate collinearity between total Met SASA 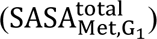 and maximum Met SASA 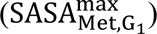 (Variance Inflation Factor = s2.4, Pearson’s *r* = 0.77, *p* = 0.0005; **Table S8**). We then evaluated both variables individually using simple linear regression analysis. As a result, total Met SASA yielded better performance, accounting for 70% of the variance in the chlorine inactivation rate constants of the studied viral strains (R^2^ = 0.70; **Table S8**). Future studies to expand the dataset with additional data points will be essential to determine whether the biological contributions of these two Met accessibility variables are distinct or inherently dependent.

Importantly, because the SASA features were identified from a broad range of virus families, our findings would have broad implications for diverse viruses. However, it is worth noting that our dataset may be constrained in capturing statistically significant contributions from other chlorine-reactive amino acids to virus inactivation if those traits are highly family-dependent. For example, Lys residues in the MS2 capsid react with chlorine to form Lys-chloramine, which subsequently reacts with Tyr residues and forms chlorotyrosine;^9^ yet this product is not observed in the PhiX174 capsid.^5^ This reaction pathway may be favored in MS2, possibly due to the limited accessibility of highly reactive residues, as the MS2 capsid contains more buried Met and entirely lacks His compared to PhiX174 capsid (**Table S5**). Furthermore, the maximum SASA of Tyr in MS2 is 2.3-fold greater than in PhiX174 (**Table S5**), suggesting that specific Tyr sites in MS2 are more accessible for reaction. Beyond Tyr, peptide-bound Lys and Arg can form chloramine rapidly when reacted with chlorine at a molar ratio of 1:1 over short timescales (< 30 min).^38^ The conversion of these positively charged side chains into neutral organic chloramines likely induces protein structural changes by disrupting critical electrostatic interactions,^39, 40^ although such mechanisms have not yet been explicitly explored in the context of virus inactivation. Nevertheless, our work represents an initial attempt to systematically evaluate virus inactivation kinetics by integrating protein folding, structural spatial arrangement, and biological function.

### Comparison of total methionine accessibility across curated structures

Viruses associated with the majority of structures in our dataset (93.4%) have not been previously characterized for their chlorine inactivation kinetics. To address this knowledge gap, we grouped the structures by virus family and then ranked them based on their median total Met SASA in the viral attachment and entry proteins 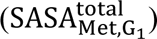, aiming to identify less-characterized viruses that are potentially insensitive to chlorine inactivation. Because this SASA value was restricted to viral attachment and entry proteins, icosahedral structures lacking these specific functional components were not included, such as the head capsids of tailed phages.^41^

At the virus family level, the total Met SASA demonstrates variations of at least six orders of magnitude (**Figure 3**). Notably, several plant viruses (*Pamosaviridae*, *Tomosaviridae*, *Bromoviridae*, *Tonesaviridae*, *Tombusviridae*, and *Tymoviridae*), and the protist virus family *Marnaviridae* were associated with relatively low median total Met SASA, ranging from 10^3^ to 10^4^ Å^2^ (**Figure 3**). Previous studies have reported that RNA sequences from *Bromoviridae*, *Marnaviridae*, *Tombusviridae*, and *Tymoviridae* remain detectable in both raw and treated wastewater.^42, 43^ Furthermore, infectious plant viruses from these groups, such as cucumber mosaic virus (*Bromoviridae*), tomato bushy stunt virus (*Tombusviridae*), and tobacco necrosis virus (*Tombusviridae*), have been isolated from surface waters.^44^ These lines of evidence confirm their potential environmental persistence. While plant and protist viruses do not pose a direct threat to human health, their high prevalence and environmental stability raise concerns for ecosystem dynamics in hydroponic agriculture and aquaculture.^45^

**Figure 3.**
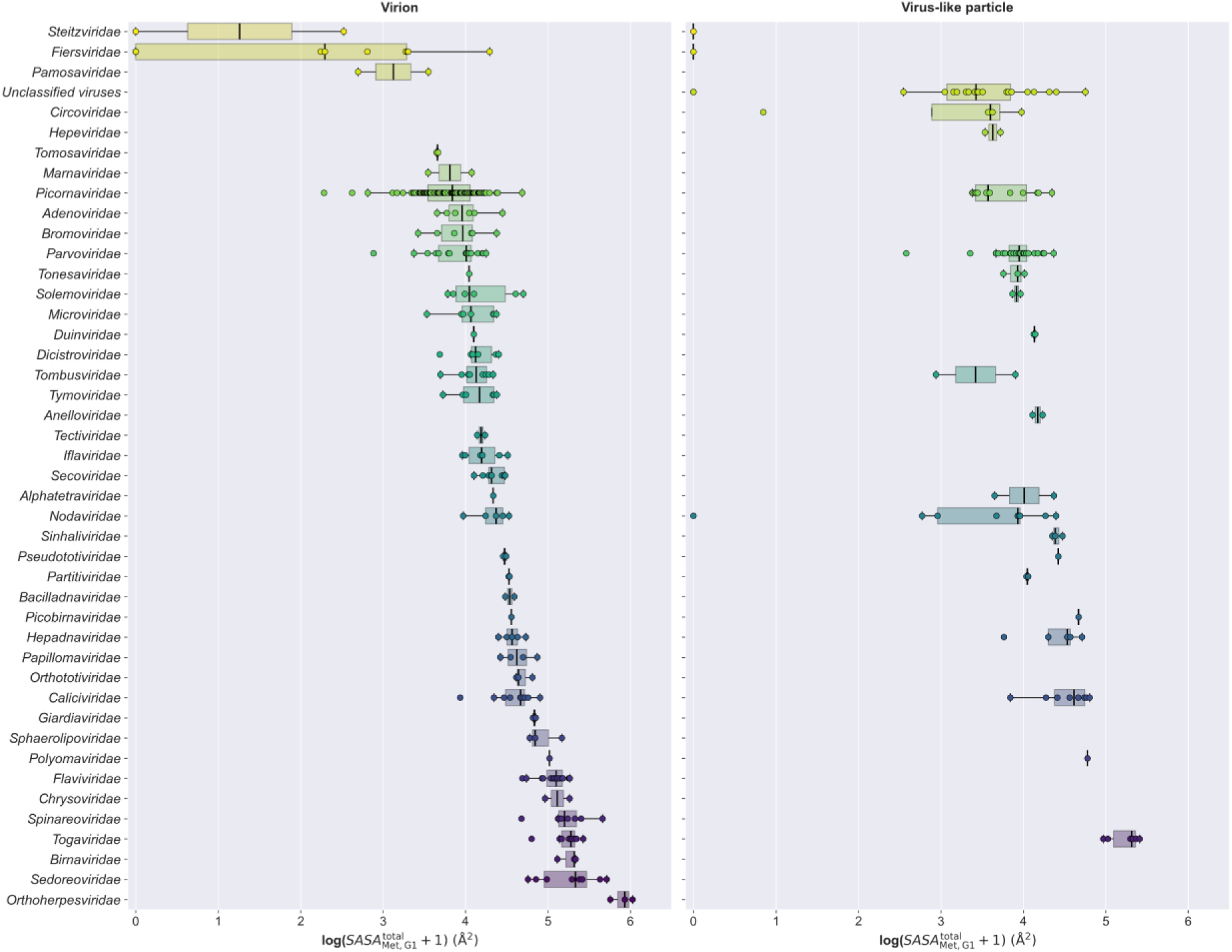
Comparison of total solvent accessible surface areas of methionine residues in the viral attachment and entry protein group 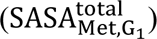 across the curated dataset. The 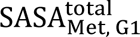 values were log-transformed. Virus families containing more than two structures were plotted and ranked by the median 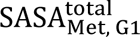 value derived from virion structures or from virus-like particle structures when virion structures were unavailable.

Beyond plant pathogens, our model identified several mammalian viruses whose attachment and entry proteins associated with remarkably low total Met SASA values (<10^3^ Å2; **Figure 3**; **Table S3**). These structures included a bat-associated circovirus (*Circoviridae*, PDB ID: 6RPO), an ssDNA virus shed in the feces of infected bats.^46^ Additionally, senecavirus A (also known as Seneca Valley virus; 6ADT and 3CJI) and Aichi virus (5AOO) of family *Picornaviridae* fell within this low total Met SASA range. Senecavirus A is an emerging pathogen that causes vesicular diseases in pigs and is transmitted primarily via direct contact and fecal contamination.^47^ While zoonotic transmission has not been reported, senecavirus A can effectively propagate in human cell lines.^48^ By comparison, Aichi viruses are human pathogens commonly co-detected with other enteric viruses in gastroenteritis patients and wastewater.^49^ A prior disinfection study on stainless steel surfaces showed that Aichi virus were more chlorine resistant compared to feline calicivirus.^50^

The total Met SASA values provide a rapid screening tool to evaluate potential chlorine resistance based solely on methionine accessibility, regardless of the inactivation contributions from reactions with other residues. To achieve a more precise estimation of virus inactivation kinetics, future modeling frameworks should integrate residue accessibility with genome reactivity alongside the efficiency of chlorine penetration across the capsid.^51^

### Comparison of SASA profiles of chlorine-reactive residues between common virus surrogates and human viruses

In studies of environmental fate, transport, and disinfection treatment, phages MS2, Q-beta (*Fiersviridae*), and PhiX174 (*Microviridae*) are widely employed as surrogates for enteroviruses (*Caliciviridae*, *Parvoviridae*, and *Picornaviridae*).^52, 53^ Likewise, phages PR772 and PRD 1 (*Tectiviridae*) serve as surrogates for adenoviruses (*Adenoviridae*),^54^ and phage Phi6 (*Cystoviridae*) is used to model enveloped human viruses.^55, 56^ The selection of surrogates for specific human viruses primarily relied on genome type similarities, envelope status, and viral particle size, yet the surface exposure of reactive amino acids in viral proteins has been largely overlooked.^57^ We then asked if the total and maximum SASA profiles of all seven chlorine-reactive residues in viral attachment and entry proteins (G1) could be applied to evaluate chlorine resistance phenotypes between virus surrogates and human viruses. To do this, we applied fuzzy c-means clustering to classify the viral structures based on their total and maximum SASA profiles.

#### Non-enveloped surrogates

All *Microviridae* structures were grouped in the same cluster as *Picornaviridae* and *Parvoviridae* for both total and maximum SASA profiles (**Figure 4**). Indeed, the total and maximum SASA profiles from these three families consistently showed relatively more buried Cys and Trp residues compared to Met and His residues, following the exposure levels of His>Met>Trp>Cys (**Figure S5 and S6**). While the *Fiersviridae* structures were clustered with *Picornaviridae* on their total SASA profiles (**Figure 4A**), none of the *Fiersviridae* structures were grouped with *Picornaviridae* on the maximum SASA profiles (**Figure 4B**). The amino acid composition of capsids from *Fiersviridae* was distinct. For example, the MS2 capsid proteins (2MS2, 9Q1D) lack His, the Q-beta capsid proteins (7LHD, 5VLY, 5VLZ, 1QBE, 5KIP) lack Met and His, and the GA capsid protein (1GAV) lacks Met, Cys, and His (**Table S3**). The lack of chlorine-reactive amino acids in the capsid could facilitate chlorine penetration and reaction with the encapsidated genome.^51^ In terms of adenovirus surrogates, PR772 (6Q5U) clustered with *Adenoviridae* on both total and maximum SASA profiles, a similarity not shared by PRD1 (1W8X). Interestingly, while the genomes of PR772 and PRD1 share greater than 97% sequence identity,^58^ the total and maximum residue SASA values of PRD1 were higher than those of PR772 for the majority of the chlorine-reactive residues (**Figure S5 and S6**); these features are consistent with the observed higher chlorine susceptibility of PRD1 compared to PR772 in previous studies (**Table S5**).

**Figure 4.**
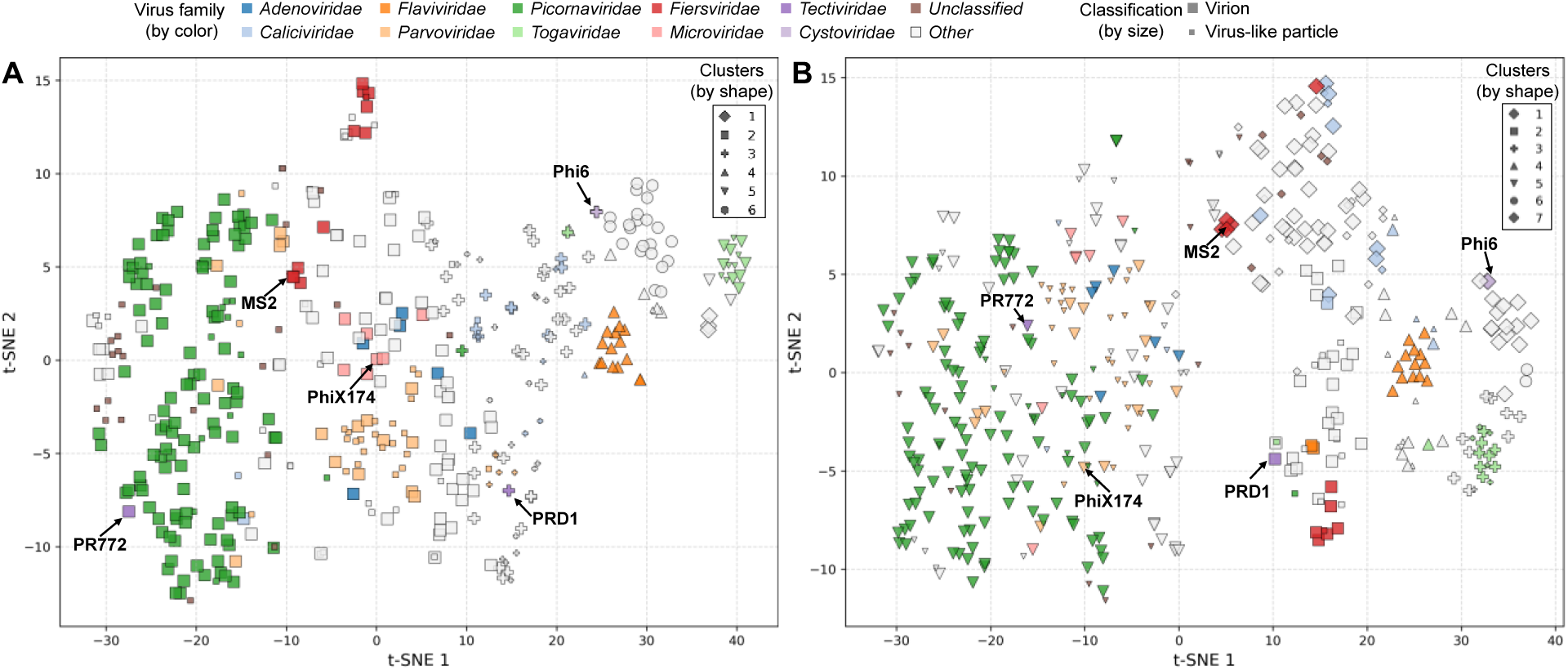
t-distributed Stochastic Neighbor Embedding (t-SNE) visualization of the fuzzy c-means clustering of viral structures based on (A) total solvent accessible surface areas (SASA) and (B) maximum SASA profiles for seven chlorine-reactive residues on essential proteins. Colors denote distinct virus families. Symbol sizes differentiate between virion and virus-like particle structures. Symbol shapes represent the specific cluster determined by the fuzzy c-means algorithm.

#### Enveloped surrogates

*Cystoviridae* phage Phi6 contains an icosahedral capsid, but its capsid SASA profiles separated from those of icosahedral capsids in enveloped human viruses belonging to *Flaviviridae* (*e.g.*, Zika and Dengue virus) and *Togaviridae* (*e.g.*, chikungunya virus) (**Figure 4**). The Phi6 capsid contains much lower total SASA values for His and Trp residues compared to *Flaviviridae* and *Togaviridae* (**Figure S5**). Notably, our dataset is focused on icosahedral architectures, we could not compare Phi6 with pleomorphic enveloped viruses, such as coronaviruses and influenza viruses, which lack icosahedral capsids.

#### Alternative surrogates

The structural clustering offers an approach for discovering alternative surrogates to cover a broader range of virus families. We found that virion and VLP structures from the same virus family consistently clustered together and were closely positioned in the t-SNE plot (**Figure 4**), indicating the structural validity of using VLPs as surrogates for treatment processes that primarily act through reactions between oxidants and proteins.^59^ VLPs have been extensively utilized in vaccine research to understand how genetic mutations of specific protein domains influence the persistence and immune responses of vaccine strains.^60, 61^ Additionally, we noted that *Picornaviridae* structures clustered more closely with several unclassified non-tailed phage structures than with the *Microviridae* structures on both total SASA- or maximum SASA-derived features (**Figure 4**). These non-tailed phage structures were VLPs constructed from metagenomic sequences of environmental samples.^62^ Future studies can use high-throughput phage isolation and genomic sequencing to discover and characterize infectious non-tailed phages,^63^ and apply them as novel, structurally validated surrogates.

### Environmental implications

Our study systematically reviewed and curated a dataset of experimentally derived, intact, environmentally relevant icosahedral virion and VLP structures from the PDB. By cross-referencing residue accessibility of these structures with chlorine inactivation rate constants from previous studies, we have demonstrated that the total and maximum SASA of Met residues in viral attachment and entry proteins significantly correlated with median virus inactivation rate constants. Furthermore, clustering based on the SASA profiles of chlorine-reactive residues effectively identifies structural similarities among capsids, which highlights the need for novel surrogates to reflect the structural diversity of human viruses. Expanding beyond chlorine-reactive residues, we quantified the total and maximum SASA for all 20 standard amino acids across each protein in our curated structures (**Table S3**). These comprehensive values can be leveraged to explore how capsid structures influence viral resistance to other protein-reactive disinfectants, such as chlorine dioxide, chloramine, and singlet oxygen.^64, 65^

While our study focused exclusively on icosahedral capsids, the developed framework is highly adaptable. Future research could expand this effort into pleomorphic enveloped viruses by incorporating the SASA of their membrane proteins responsible for attachment. Several limitations of this study should be noted. First, virus inactivation by free chlorine can be driven by multiple mechanisms. We only evaluated resistance from the protein structural perspective. Second, our analysis was restricted to clean water and buffered systems to maintain consistency with the experimental conditions under which the PDB structures were resolved. Because complex water matrices can significantly alter capsid integrity, future studies need to evaluate the influence of environmental water chemistry on virus capsid stability to expand our framework to broader treatment conditions.

## ASSOCIATED CONTENT

### Supporting information

The supporting information is available free of charge.

Figures of flowchart of systematic retrieval, screening, and selection of icosahedral viral capsid structures, Phi6 inactivation kinetics by free chlorine, comparative analysis of residue SASA calculated using various parameters, total SASA of chlorine-reactive residues in viral attachment and entry proteins G1 across the selected viral strains, total SASA profiles of chlorine-reactive residues in viral attachment and entry proteins G1 across virion and VLP structures in the curated dataset, and maximum SASA profiles of chlorine-reactive residues in viral attachment and entry proteins G1 across virion and VLP structures in the curated dataset (PDF).

Tables of the curated dataset, summary of genomic, protein, and structural characteristics of virus families within the curated dataset, comprehensive total and maximum SASA of 20 standard amino acids for each viral protein in the curated dataset, structures and parameters used in the systematic evaluation of SASA workflow, chlorine inactivation rate constants and SASA features of selected viral strains, Pearson correlation analysis of SASA values calculated using various parameters, Spearman and Kendall correlation analyses between *k_obs_* and SASA matrices for the selected viral strains, multiple linear regression analysis, and fuzzy c-means clustering (XLSX).

## ACKNOWLEDGEMENTS

This study was funded by the National Science Foundation (CBET 2212779) and the University at Buffalo Research and Economic Development Grant Program to Y.Y., and the National Institutes of Health Human Virome Program (U01DE035632) to K.K. and Y.Y.. C.Z. was supported by a graduate assistantship from the School of Engineering and Applied Sciences at the University at Buffalo. K.P. was the recipient of the Environmental Engineering and Science Foundation Scholarship. S.T. was supported by the Japan Society for the Promotion of Science Grant-in-Aid for Scientific Research B (JP26K01072).

## References

(1) Chaplin, M.; Leung, K.; Szczuka, A.; Hansen, B.; Rockey, N. C.; Henderson, J. B.; Wigginton, K. R. Linear Mixed Model of Virus Disinfection by Free Chlorine to Harmonize Data Collected across Broad Environmental Conditions. Environmental science & technology 2024, 58 (27), 12260–12271.

(2) Terpstra, F. G.; van den Blink, A. E.; Bos, L. M.; Boots, A. G.; Brinkhuis, F. H.; Gijsen, E.; van Remmerden, Y.; Schuitemaker, H.; van ‘t Wout, A. B. Resistance of surface-dried virus to common disinfection procedures. J Hosp Infect 2007, 66 (4), 332–338.

(3) Mayer, B. K.; Yang, Y.; Gerrity, D. W.; Abbaszadegan, M. The Impact of Capsid Proteins on Virus Removal and Inactivation During Water Treatment Processes. Microbiology insights 2015, 8 (2), 15–28.

(4) Sano, D.; Pintó, R. M.; Omura, T.; Bosch, A. Detection of Oxidative Damages on Viral Capsid Protein for Evaluating Structural Integrity and Infectivity of Human Norovirus. Environmental Science & Technology 2010, 44 (2), 808–812.

(5) Zhu, C.; Ye, Y. Reactivity of Viral Proteins with Free Chlorine: Structural Insights and Implications for Virus Inactivation. Environmental Science & Technology 2025, 59 (32), 17188–17197.

(6) Ye, Y.; Chang, P. H.; Hartert, J.; Wigginton, K. R. Reactivity of Enveloped Virus Genome, Proteins, and Lipids with Free Chlorine and UV254. Environmental Science & Technology 2018, 52 (14), 7698–7708.

(7) Wigginton, K. R.; Pecson, B. M.; Sigstam, T.; Bosshard, F.; Kohn, T. Virus inactivation mechanisms: impact of disinfectants on virus function and structural integrity. Environ Sci Technol 2012, 46 (21), 12069–12078.

(8) Pattison, D. I.; Davies, M. J. Absolute Rate Constants for the Reaction of Hypochlorous Acid with Protein Side Chains and Peptide Bonds. Chemical Research in Toxicology 2001, 14 (10), 1453–1464.

(9) Choe, J. K.; Richards, D. H.; Wilson, C. J.; Mitch, W. A. Degradation of Amino Acids and Structure in Model Proteins and Bacteriophage MS2 by Chlorine, Bromine, and Ozone. Environmental Science & Technology 2015, 49 (22), 13331–13339.

(10) Sigstam, T.; Gannon, G.; Cascella, M.; Pecson, B. M.; Wigginton, K. R.; Kohn, T. Subtle Differences in Virus Composition Affect Disinfection Kinetics and Mechanisms. Appl Environ Microbiol 2013, 79 (11), 3455–3467.

(11) Suckau, D.; Mak, M.; Przybylski, M. Protein surface topology-probing by selective chemical modification and mass spectrometric peptide mapping. Proceedings of the National Academy of Sciences 1992, 89 (12), 5630–5634.

(12) Lundeen, R. A.; McNeill, K. Reactivity Differences of Combined and Free Amino Acids: Quantifying the Relationship between Three-Dimensional Protein Structure and Singlet Oxygen Reaction Rates. Environmental Science & Technology 2013, 47 (24), 14215–14223.

(13) Johnson, J. E.; Olson, A. J. Icosahedral virus structures and the protein data bank. J Biol Chem 2021, 296, 100554.

(14) Berman, H. M.; Westbrook, J.; Feng, Z.; Gilliland, G.; Bhat, T. N.; Weissig, H.; Shindyalov, I. N.; Bourne, P. E. The Protein Data Bank. Nucleic Acids Research 2000, 28 (1), 235–242.

(15) Rose, Y.; Duarte, J. M.; Lowe, R.; Segura, J.; Bi, C.; Bhikadiya, C.; Chen, L.; Rose, A. S.; Bittrich, S.; Burley, S. K.;, et al. RCSB Protein Data Bank: Architectural Advances Towards Integrated Searching and Efficient Access to Macromolecular Structure Data from the PDB Archive. Journal of Molecular Biology 2021, 433 (11), 166704.

(16) Ahmad, S.; Jose da Costa Gonzales, L.; Bowler-Barnett, Emily H.; Rice, Daniel L.; Kim, M.; Wijerathne, S.; Luciani, A.; Kandasaamy, S.; Luo, J.; Watkins, X.; et al. The UniProt website API: facilitating programmatic access to protein knowledge. Nucleic Acids Research 2025, 53 (W1), W547–W553.

(17) Lefkowitz, E.; Smith, D. B.; Hendrickson, R.; International Committee on Taxonomy of Viruses. ICTV Virus Metadata Resource (VMR) [VMR_MSL40.v3.20260223]. 2026. https://ictv.global/vmr (accessed 2026-03-01).

(18) Mihara, T.; Nishimura, Y.; Shimizu, Y.; Nishiyama, H.; Yoshikawa, G.; Uehara, H.; Hingamp, P.; Goto, S.; Ogata, H. Linking Virus Genomes with Host Taxonomy. Viruses 2016, 8 (3), 66.

(19) De Castro, E.; Hulo, C.; Masson, P.; Auchincloss, A.; Bridge, A.; Le Mercier, P. ViralZone 2024 provides higher-resolution images and advanced virus-specific resources. Nucleic Acids Research 2024, 52 (D1), D817–D821.

(20) PDB Archive biological assembly cordinate files in mmCIF format. 2023. https://files.wwpdb.org/pub/pdb/data/assemblies/mmCIF (accessed 2026-05-18).

(21) Cock, P. J. A.; Antao, T.; Chang, J. T.; Chapman, B. A.; Cox, C. J.; Dalke, A.; Friedberg, I.; Hamelryck, T.; Kauff, F.; Wilczynski, B.;, et al. Biopython: freely available Python tools for computational molecular biology and bioinformatics. Bioinformatics 2009, 25 (11), 1422–1423.

(22) Schrödinger, L. The PyMOL Molecular Graphics System (Version 3.1.7.2). 2025. https://pymol.org/conda/ (accessed.

(23) Shrake, A.; Rupley, J. A. Environment and exposure to solvent of protein atoms. Lysozyme and insulin. Journal of Molecular Biology 1973, 79 (2), 351–371.

(24) Torii, S.; Gouttenoire, J.; Kumar, K.; Antanasijevic, A.; Kohn, T. Influence of Amino Acid Substitutions in Capsid Proteins of Coxsackievirus B5 on Free Chlorine and Thermal Inactivation. Environmental Science & Technology 2024, 58 (12), 5279–5289.

(25) O’brien, R. M. A Caution Regarding Rules of Thumb for Variance Inflation Factors. Qual Quant 2007, 41 (5), 673–690.

(26) Bezdek, J. C.; Ehrlich, R.; Full, W. FCM: The fuzzy c-means clustering algorithm. Computers & Geosciences 1984, 10 (2-3), 191–203.

(27) Maaten, L. v. d.; Hinton, G. E. Visualizing Data using t-SNE. Journal of Machine Learning Research 2008, 9, 2579–2605.

(28) Zeltins, A. Construction and Characterization of Virus-Like Particles: A Review. Molecular Biotechnology 2013, 53 (1), 92–107.

(29) Li, J.; Predmore, A.; Divers, E.; Lou, F. New Interventions Against Human Norovirus: Progress, Opportunities, and Challenges. Annual Review of Food Science and Technology 2012, 3 (Volume 3, 2012), 331–352.

(30) Baltimore, D. Expression of animal virus genomes. Bacteriological Reviews 1971, 35 (3), 235–241.

(31) Noda, T. Native Morphology of Influenza Virions. Front. Microbio. 2012, 2.

(32) Neuman, B. W.; Adair, B. D.; Yoshioka, C.; Quispe, J. D.; Orca, G.; Kuhn, P.; Milligan, R. A.; Yeager, M.; Buchmeier, M. J. Supramolecular Architecture of Severe Acute Respiratory Syndrome Coronavirus Revealed by Electron Cryomicroscopy. J Virol 2006, 80 (16), 7918–7928.

(33) Sivey, J. D.; Roberts, A. L. Assessing the Reactivity of Free Chlorine Constituents Cl2, Cl2O, and HOCl Toward Aromatic Ethers. Environmental Science & Technology 2012, 46 (4), 2141–2147.

(34) Wang, J.; Krudy, G.; Hou, T.; Zhang, W.; Holland, G.; Xu, X. Development of Reliable Aqueous Solubility Models and Their Application in Druglike Analysis. Journal of Chemical Information and Modeling 2007, 47 (4), 1395–1404.

(35) Mitternacht, S. FreeSASA: An open source C library for solvent accessible surface area calculations. F1000Research 2016, 5 (189).

(36) Miseta, A.; Csutora, P. Relationship Between the Occurrence of Cysteine in Proteins and the Complexity of Organisms. Molecular Biology and Evolution 2000, 17 (8), 1232–1239.

(37) Nagano, N.; Ota, M.; Nishikawa, K. Strong hydrophobic nature of cysteine residues in proteins. FEBS Lett 1999, 458 (1), 69–71.

(38) Shi, J. L.; Mitch, W. A. Lysine and Arginine Reactivity and Transformation Products during Peptide Chlorination. Environmental Science & Technology 2023, 57 (14), 5852–5860.

(39) Müller, A.; Langklotz, S.; Lupilova, N.; Kuhlmann, K.; Bandow, J. E.; Leichert, L. I. O. Activation of RidA chaperone function by N-chlorination. Nat Commun 2014, 5 (1), 5804.

(40) Varatnitskaya, M.; Fasel, J.; Müller, A.; Lupilov, N.; Shi, Y.; Fuchs, K.; Krewing, M.; Jung, C.; Jacob, T.; Sitek, B.;, et al. An increase in surface hydrophobicity mediates chaperone activity in N-chlorinated RidA. Redox Biology 2022, 53, 102332.

(41) Nobrega, F. L.; Vlot, M.; De Jonge, P. A.; Dreesens, L. L.; Beaumont, H. J. E.; Lavigne, R.; Dutilh, B. E.; Brouns, S. J. J. Targeting mechanisms of tailed bacteriophages. Nat Rev Microbiol 2018, 16 (12), 760–773.

(42) Bačnik, K.; Kutnjak, D.; Pecman, A.; Mehle, N.; Tušek Žnidarič, M.; Gutiérrez Aguirre, I.; Ravnikar, M. Viromics and infectivity analysis reveal the release of infective plant viruses from wastewater into the environment. Water Research 2020, 177, 115628.

(43) Segelhurst, E.; Bard, J. E.; Gallo, S.; Huang, V.; Pohlman, A.; Yergeau, D. A.; Surtees, J. A.; Bradley, I. M.; Ye, Y. Recovering wastewater RNA for virome sequencing by systematically optimized tangential-flow ultrafiltration and Nanotrap microbiome particles. Appl Environ Microbiol 2025, 91 (10), e00777–00725.

(44) Mehle, N.; Ravnikar, M. Plant viruses in aqueous environment – Survival, water mediated transmission and detection. Water Research 2012, 46 (16), 4902–4917.

(45) Park, W. M.; Lee, G. P.; Ryu, K. H.; Park, K. W. Transmission of tobacco mosaic virus in recirculating hydroponic system. Scientia Horticulturae 1999, 79 (3-4), 217–226.

(46) Lecis, R.; Mucedda, M.; Pidinchedda, E.; Zobba, R.; Pittau, M.; Alberti, A. Genomic characterization of a novel bat-associated Circovirus detected in European Miniopterus schreibersii bats. Virus Genes 2020, 56 (3), 325–328.

(47) Wu, H.; Li, C.; Ji, Y.; Mou, C.; Chen, Z.; Zhao, J. The Evolution and Global Spatiotemporal Dynamics of Senecavirus A. Microbiol Spectr 2022, 10 (6), e0209022.

(48) Yang, M.; van Bruggen, R.; Xu, W. Generation and diagnostic application of monoclonal antibodies against Seneca Valley virus. J Vet Diagn Invest 2012, 24 (1), 42–50.

(49) Kitajima, M.; Gerba, C. P. Aichi virus 1: environmental occurrence and behavior. Pathogens 2015, 4 (2), 256–268.

(50) Cromeans, T.; Park, G. W.; Costantini, V.; Lee, D.; Wang, Q.; Farkas, T.; Lee, A.; Vinje, J. Comprehensive comparison of cultivable norovirus surrogates in response to different inactivation and disinfection treatments. Appl Environ Microbiol 2014, 80 (18), 5743–5751.

(51) Qiao, Z.; Ye, Y.; Szczuka, A.; Harrison, K. R.; Dodd, M. C.; Wigginton, K. R. Reactivity of Viral Nucleic Acids with Chlorine and the Impact of Virus Encapsidation. Environ Sci Technol 2022, 56 (1), 218–227.

(52) Kuzmanovic, D. A.; Elashvili I Fau - Wick, C.; Wick C Fau - O’Connell, C.; O’Connell C Fau - Krueger, S.; Krueger, S. Bacteriophage MS2: molecular weight and spatial distribution of the protein and RNA components by small-angle neutron scattering and virus counting. 2003, (0969-2126 (Print)).

(53) Gerba, C. P.; Abd-Elmaksoud, S.; Newick, H.; El-Esnawy, N. A.; Barakat, A.; Ghanem, H. Assessment of Coliphage Surrogates for Testing Drinking Water Treatment Devices. Food Environ Virol 2015, 7 (1), 27–31.

(54) Gall, A. M.; Shisler, J. L.; Mariñas, B. J. Characterizing Bacteriophage PR772 as a Potential Surrogate for Adenovirus in Water Disinfection: A Comparative Analysis of Inactivation Kinetics and Replication Cycle Inhibition by Free Chlorine. Environmental Science & Technology 2016, 50 (5), 2522–2529.

(55) Aquino De Carvalho, N.; Stachler, E. N.; Cimabue, N.; Bibby, K. Evaluation of Phi6 Persistence and Suitability as an Enveloped Virus Surrogate. Environmental Science & Technology 2017, 51 (15), 8692–8700.

(56) Ye, Y.; Ellenberg, R. M.; Graham, K. E.; Wigginton, K. R. Survivability, Partitioning, and Recovery of Enveloped Viruses in Untreated Municipal Wastewater. Environmental Science & Technology 2016, 50 (10), 5077–5085.

(57) Sinclair, R. G.; Rose, J. B.; Hashsham, S. A.; Gerba, C. P.; Haas, C. N. Criteria for Selection of Surrogates Used To Study the Fate and Control of Pathogens in the Environment. Appl Environ Microbiol 2012, 78 (6), 1969–1977.

(58) Lute, S.; Aranha, H.; Tremblay, D.; Liang, D.; Ackermann, H. W.; Chu, B.; Moineau, S.; Brorson, K. Characterization of coliphage PR772 and evaluation of its use for virus filter performance testing. Appl Environ Microbiol 2004, 70 (8), 4864–4871.

(59) Caballero, S.; Abad, F. X.; Loisy, F.; Le Guyader, F. S.; Cohen, J.; Pintó, R. M.; Bosch, A. Rotavirus Virus-Like Particles as Surrogates in Environmental Persistence and Inactivation Studies. Appl Environ Microbiol 2004, 70 (7), 3904–3909.

(60) Hankaniemi, M. M.; Baikoghli, M. A.; Stone, V. M.; Xing, L.; Väätäinen, O.; Soppela, S.; Sioofy-Khojine, A.; Saarinen, N. V. V.; Ou, T.; Anson, B.;, et al. Structural Insight into CVB3-VLP Non-Adjuvanted Vaccine. Microorganisms 2020, 8 (9), 1287.

(61) Soppela, S.; Plavec, Z.; Gröhn, S.; Mustonen, I.; Jartti, M.; Oikarinen, S.; Laajala, M.; Marjomäki, V.; Butcher, S. J.; Hankaniemi, M. M. Immunological and structural evaluation of the intranasally administrated CVB1 whole-virus and VLP vaccines. Scientific Reports 2025, 15 (1), 10198.

(62) Rūmnieks, J.; Liekniņa, I.; Kalniņš, G.; Šišovs, M.; Akopjana, I.; Bogans, J.; Tārs, K. Three-dimensional structure of 22 uncultured ssRNA bacteriophages: Flexibility of the coat protein fold and variations in particle shapes. Science Advances 2020, 6 (36), eabc0023.

(63) Kauffman, K. M.; Hussain, F. A.; Yang, J.; Arevalo, P.; Brown, J. M.; Chang, W. K.; VanInsberghe, D.; Elsherbini, J.; Sharma, R. S.; Cutler, M. B.;, et al. A major lineage of non-tailed dsDNA viruses as unrecognized killers of marine bacteria. Nature 2018, 554 (7690), 118–122.

(64) Wigginton, K. R.; Kohn, T. Virus disinfection mechanisms: the role of virus composition, structure, and function. Curr Opin Virol 2012, 2 (1), 84–89.

(65) Di Mascio, P.; Martinez, G. R.; Miyamoto, S.; Ronsein, G. E.; Medeiros, M. H. G.; Cadet, J. Singlet Molecular Oxygen Reactions with Nucleic Acids, Lipids, and Proteins. Chem Rev 2019, 119 (3), 2043–2086.

