## Supplemental figures for "Solvent accessibility of chlorine-reactive amino acid residues in icosahedral virus structures: A meta-analysis"

---


**Summary: 10 Pages, 5 Figures, 9 Tables (all SI tables are in a separate XLSX file)**

**Table of Contents**

**Figure S1**. Flowchart of systematic retrieval, screening, and selection of icosahedral viral capsid structures for the meta-analysis.

**Figure S2.** Phi6 inactivation by free chlorine (3.1 ± 0.1 mg/L as $Cl_2$) in phosphate buffer (5 mM $PO_4^{3-}$, 10 mM NaCl, pH 7.5).

**Figure S3.** Comparative analysis of solvent accessible surface area (SASA) values per residue calculated using various calculation parameters for assembled capsid structures (PDB ID: 8AT5, 5C4W, 7BG6, and 2BPA).

**Figure S4.** Total SASA of seven chlorine-reactive residues in the group of viral attachment and entry proteins (G1) across the selected viral strains.

**Figure S5.** Total SASA profiles of seven chlorine-reactive residues in the group of viral attachment and entry proteins (G1) across virion and virus-like particle (VLP) structures in the curated dataset.

**Figure S6.** Maximum SASA profiles of seven chlorine-reactive residues in the group of viral attachment and entry proteins (G1) across virion and virus-like particle (VLP) structures in the curated dataset.

**Table S1.** Curated dataset of icosahedral virus structures from the Protein Data Bank.

**Table S2.** Summary of genomic, protein, and structural characteristics of virus families within the curated dataset

**Table S3.** Comprehensive total and maximum SASA of 20 standard amino acid residues for each viral protein in the curated dataset.

**Table S4.** Structures and parameters used in the systematic evaluation of SASA calculation workflow.

**Table S5.** Chlorine inactivation rate constants and SASA characteristics of the selected viral strains.

**Table S6.** Pearson correlation analyses of SASA values calculated using various probe radii (r, Å) and surface resolutions (n).

**Table S7.** Spearsman and Kendall correlation analyses between median chlorine inactivation rate constants ($k_{obs}$) against total and maximum residue SASA (total or maximum) across the groups of viral attachment and entry proteins (G1) and other proteins (G2) for the selected viral strains.

**Table S8.** Multiple linear regression analysis of median virus inactivation rate constant ($k_{obs}$) as a function of total and maximum SASA of Met in the group of viral attachment and entry proteins (G1).

**Table S9.** Fuzzy c-means clustering of curated PDB structure based on total or maximum SASA of chlorine-reactive residues in the group of viral attachment and entry proteins (G1).

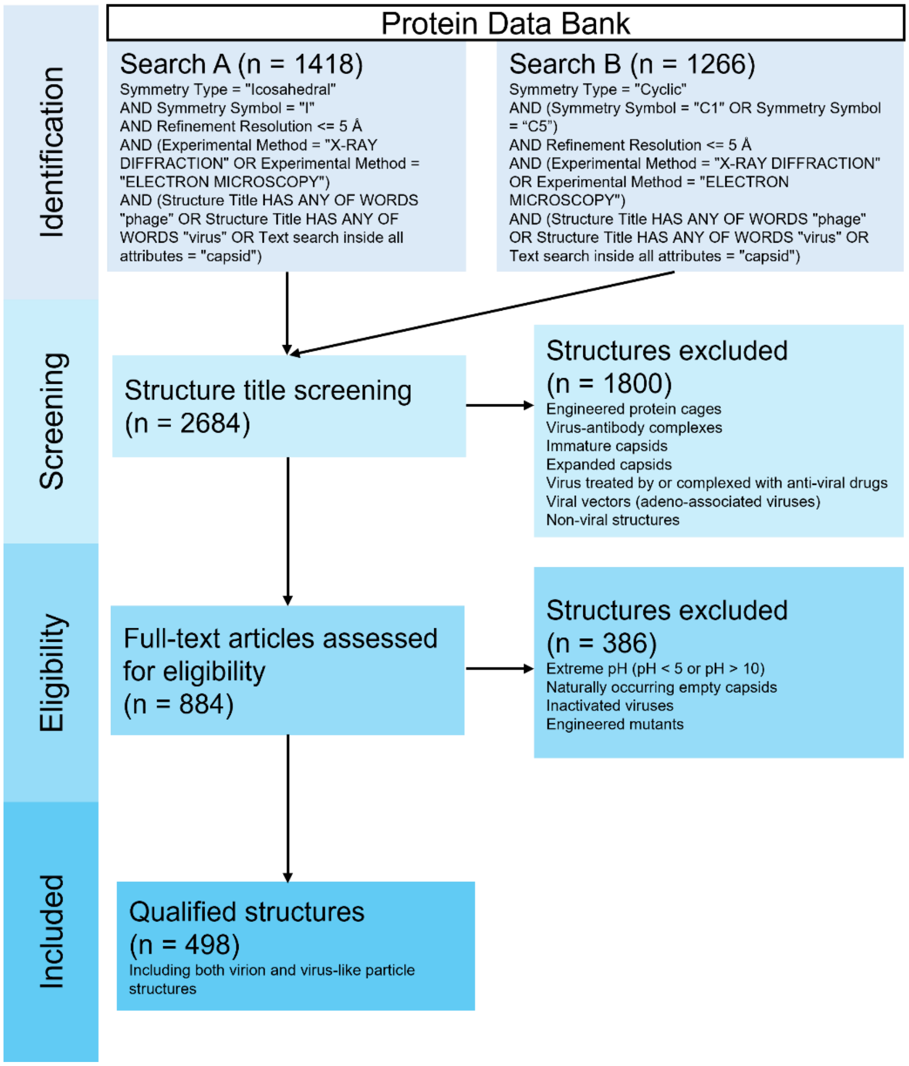


**Figure S1**. Flowchart of systematic retrieval, screening, and selection of icosahedral viral capsid structures for the meta-analysis.

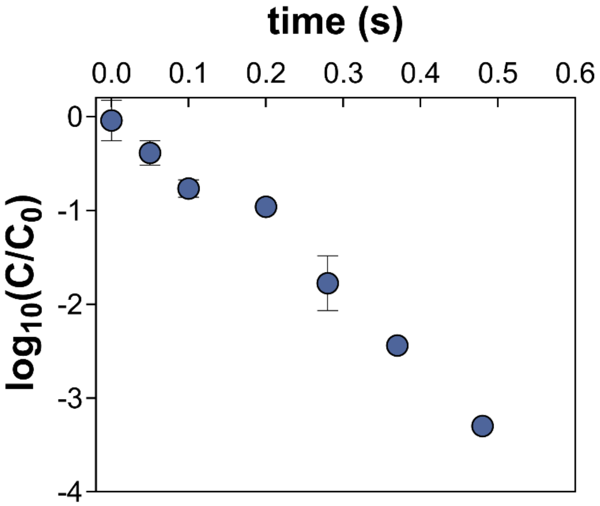


**Figure S2.** Phi6 inactivation by free chlorine (3.1 ± 0.1 mg/L as $Cl_2$) in phosphate buffer (5 mM $PO_4^{3-}$, 10 mM NaCl, pH 7.5). The reaction was performed in a continuous quench-flow reactor and quenched with 50 mM Tris-HCl as described previously.[1] Pseudo-first-order kinetics was applied to calculate the Phi6 inactivation rate constant as 281.7 (L $mg^{-1}$ $min^{-1}$).

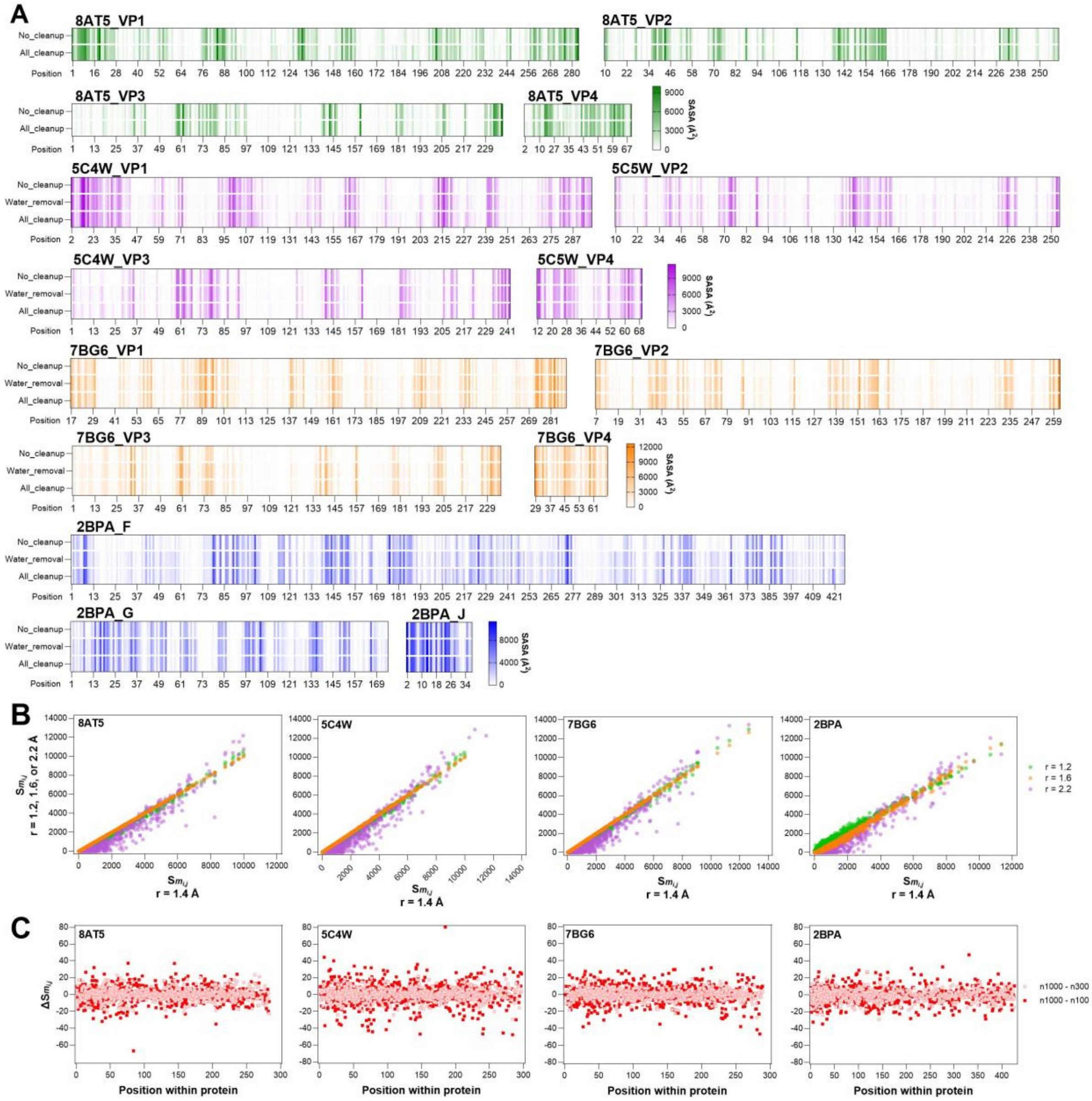


**Figure S3.** Comparative analysis of solvent accessible surface area (SASA) values per residue calculated using various calculation parameters for assembled capsid structures (PDB ID: 8AT5, 5C4W, 7BG6, and 2BPA). (A) Comparison of residue SASA values ($S_{m_{i,j}}$) calculated from the raw structural file (no cleanup), and the raw structure with the removal of water molecules (water removal) and all non-amino acid ligands and ions (all cleanup); (B) Correlations between the residue SASA values ($S_{m_{i,j}}$) calculated using probe radii of 1.2, 1.6, and 2.2 Å with those calculated using the default radius of 1.4 Å; (C) Residue SASA discrepancies ($\Delta S_{m_{i,j}}$) resulting from variations in surface resolution, specifically comparing a high-resolution baseline (n = 1000) against lower resolutions (n = 100 and 300).

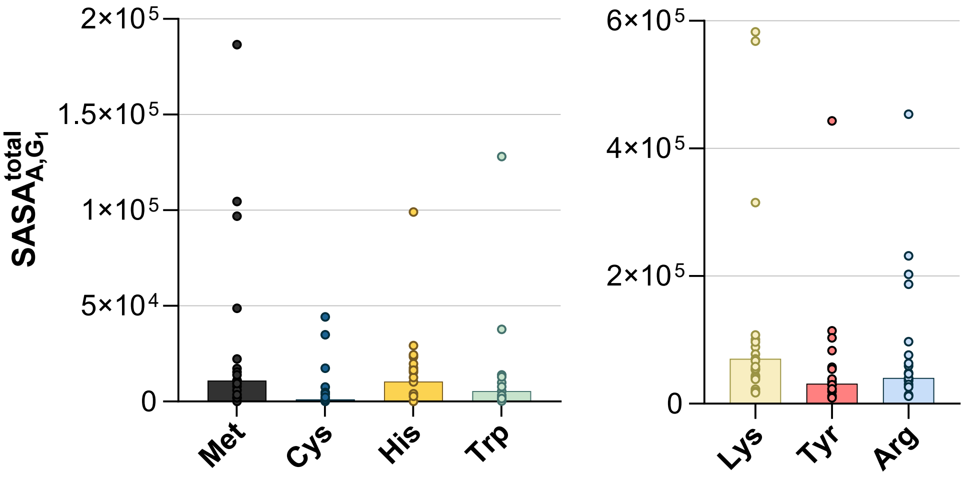


**Figure S4.** Total SASA of seven chlorine-reactive residues in the group of viral attachment and entry proteins (G1) across the selected viral strains. Bars represent the median total SASA values.

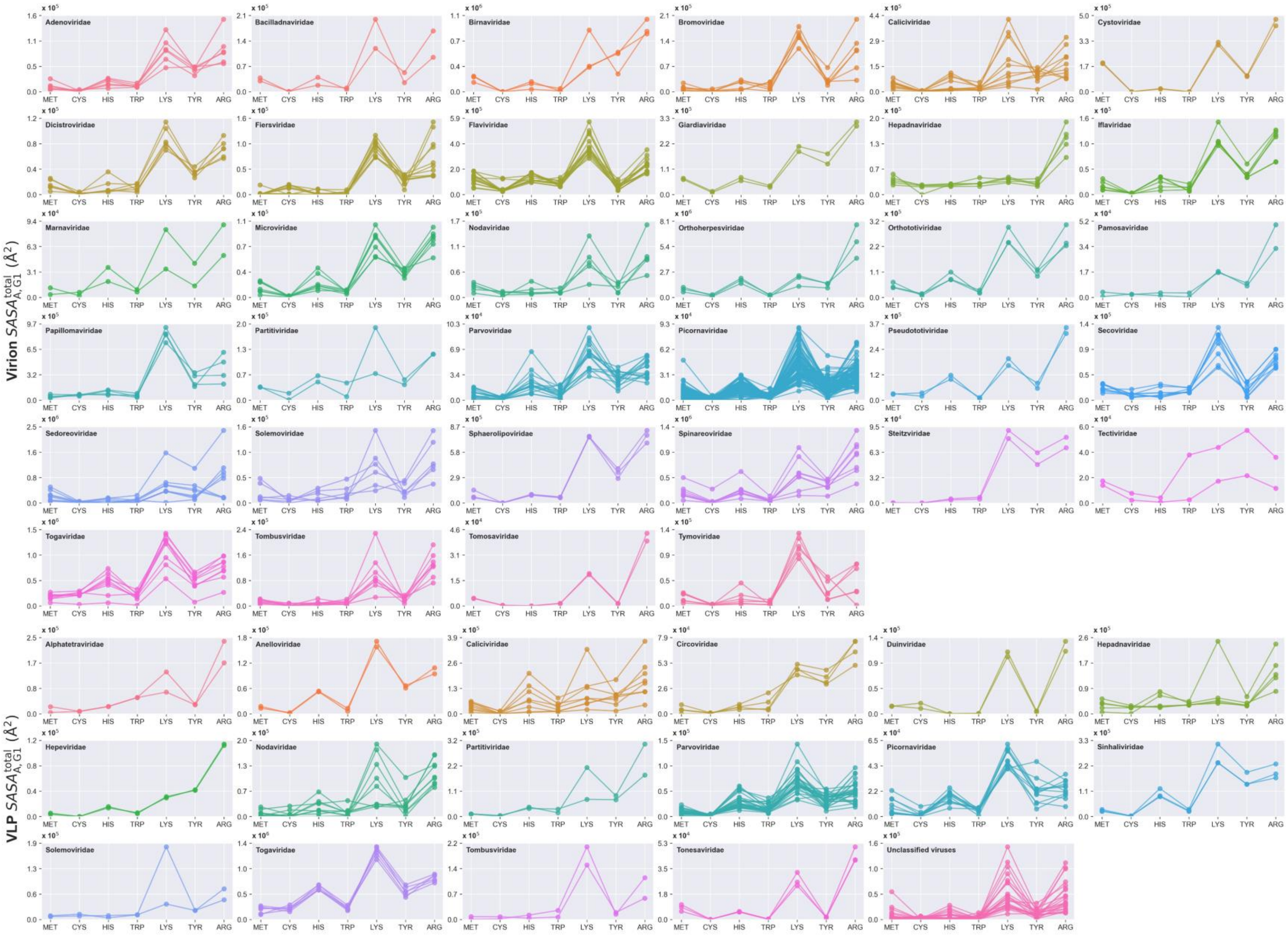
Virion $SASA^{total}_{A,G1}$ (Å$^2$)
VLP $SASA^{total}_{A,G1}$ (Å$^2$)
Adenoviridae
Bacilladnaviridae
Birnaviridae
Bromoviridae
Caliciviridae
Cystoviridae
Dicistroviridae
Fiersviridae
Flaviviridae
Giardiaviridae
Hepadnaviridae
Iflaviridae
Marnaviridae
Microviridae
Nodaviridae
Orthoherpesviridae
Orthototiviridae
Pamosaviridae
Papillomaviridae
Partitiviridae
Parvoviridae
Picornaviridae
Pseudototiviridae
Secoviridae
Sedoreoviridae
Solemoviridae
Sphaerolipoviridae
Spinareoviridae
Steitzviridae
Tectiviridae
Togaviridae
Tombusviridae
Tomosaviridae
Tymoviridae
Alphatetraviridae
Anelloviridae
Caliciviridae
Circoviridae
Duinviridae
Hepadnaviridae
Hepeviridae
Nodaviridae
Partitiviridae
Parvoviridae
Picornaviridae
Sinhaliviridae
Solemoviridae
Togaviridae
Tombusviridae
Tonesaviridae
Unclassified viruses
MET CYS HIS TRP LYS TYR ARG

**Figure S5.** Total SASA profiles of seven chlorine-reactive residues in the group of viral attachment and entry proteins (G1) across virion (top 6 rows) and virus-like particle (VLP; bottom 3 rows) structures in the curated dataset.

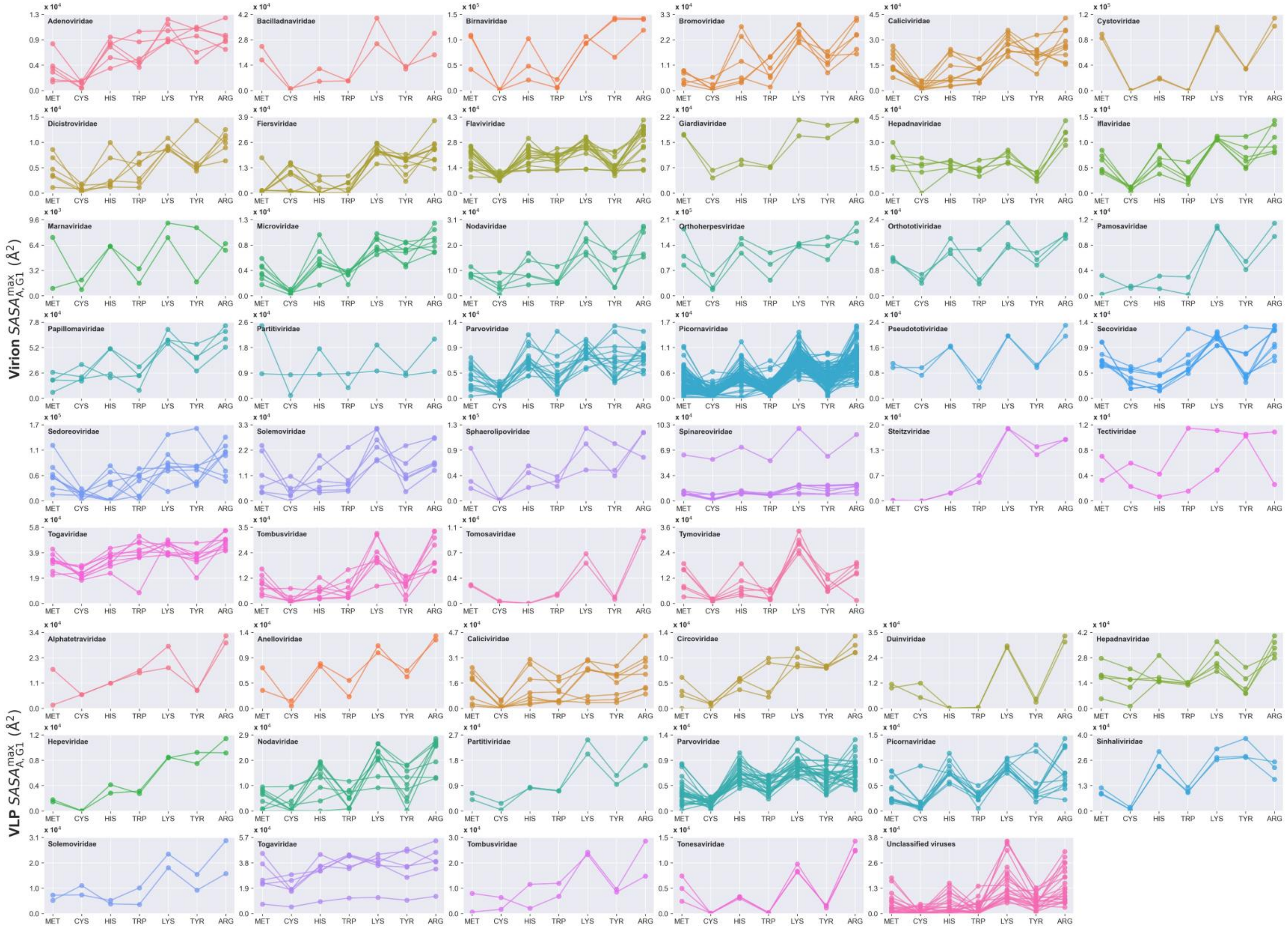
Virion $SASA_{A,G1}^{max}$ (Å$^2$)
VLP $SASA_{A,G1}^{max}$ (Å$^2$)
MET CYS HIS TRP LYS TYR ARG
Adenoviridae
Bacilladnaviridae
Birnaviridae
Bromoviridae
Caliciviridae
Cystoviridae
Dicistroviridae
Fiersviridae
Flaviviridae
Giardiaviridae
Hepadnaviridae
Iflaviridae
Marnaviridae
Microviridae
Nodaviridae
Orthoherpesviridae
Orthototiviridae
Pamosaviridae
Papillomaviridae
Partitiviridae
Parvoviridae
Picornaviridae
Pseudototiviridae
Secoviridae
Sedoreoviridae
Solemoviridae
Sphaerolipoviridae
Spinareoviridae
Steitzviridae
Tectiviridae
Togaviridae
Tombusviridae
Tomosaviridae
Tymoviridae
Alphatetraviridae
Anelloviridae
Caliciviridae
Circoviridae
Duinviridae
Hepadnaviridae
Hepeviridae
Nodaviridae
Partitiviridae
Parvoviridae
Picornaviridae
Sinhaliviridae
Solemoviridae
Togaviridae
Tombusviridae
Tonesaviridae
Unclassified viruses

**Figure S6.** Maximum SASA profiles of seven chlorine-reactive residues in the group of viral attachment and entry proteins (G1) across virion (top 6 rows) and virus-like particle (VLP; bottom 3 rows) structures in the curated dataset.

References

(1) Zhu, C.; Ye, Y. Reactivity of Viral Proteins with Free Chlorine: Structural Insights and Implications for Virus Inactivation. *Environmental Science & Technology* **2025**, *59* (32), 17188-17197
